# Contrasting urban adaptations of seed dispersal traits in two closely-related plant species

**DOI:** 10.1101/2023.12.27.573339

**Authors:** Shuhei H Nakano, Atushi Ushimaru

## Abstract

- Urbanisation has largely altered city landscapes, resulting in habitat loss, fragmentation, and environmental degradation. City plants often have adapting traits and ecologies to such urban habitat environments. The existing literature on urban adaptation has often focused on species that shrink the abundances in urban areas compared to rural areas. Meanwhile, the studies on species that expand their range in cities remain limited, although they would exhibit different trait responses to urban environments, particularly in dispersal-related traits.
- In this study, we compared seed dispersal-related traits between *Youngia japonica* subsp. *japonica (Y.j. japonica)*, which is expanding its range in urban areas and its closely-relative *Y.j. elstonii*, which is contracting its range in urban areas along an urban-rural gradient of the Osaka-Kobe megacity area. We also examined relationships between seed traits and terminal velocity which influences seed dispersal ability.
- We found that achene length increased with the degree of urbanisation only in *Y.j. japonica*, while pappus length decreased only in *Y.j. elstonii*. The terminal velocity had a significant relationship only with pappus length in *Y. japonic*a.
- Seed traits of two subspecies with different urban distributions showed different responses to urbanisation. Our results suggest that species expanding their distribution in urban areas has increased seed competitiveness while maintaining dispersal ability. In contrast, species that has shrunk in urban areas likely showed a passive response to urban fragmentation, indicating a loss of seed dispersal ability.

## INTRDUCTION

In the 21st century, global urbanisation, the concentration of populations in urban areas in the world, has largely altered ecological environments of life. The expansion of man-made surfaces in urban and peri-urban areas has resulted in habitat loss and fragmentation for organisms living in natural and semi-natural ecosystems (Dobbs et al., 2017; Pauleit et al., 2005). Additionally, it has caused environmental degradation in the remaining habitats (Pickett et al., 2001; Sukopp 2004; Grimm et al., 2008). When comparing urban areas to nearby rural landscapes in Europe and North America, fragmentation and eutrophication, increased pH levels, and water shortage in soils are conspicuous in urban green spaces (Pickett et al., 2011; Johnson et al., 2015). These changes in habitat environments have caused significant reductions in the populations of native plant and animal species, leading to local extinctions and ultimately biodiversity decline in urban areas (Williams et al., 2005; Uchida et al 2018). In contrast, current research has indicated that certain plant species can survive and exhibit unique trait adaptations to urban habitat settings (Johnson & Munshi-South, 2017; Rivkin et al., 2019; Lembert et al., 2021).

Comparison of populations between urban and nearby rural landscapes have revealed prevalent patterns of vegetative and reproductive traits of multiple plant species within urban settings. These observations imply the presence of similar trait variations among species in reaction to environmental shifts linked to urbanisation (Williams et al., 2015; Cochard et al., 2019). For example, various species have been reported to exhibit increased plant height owing to environmental changes such as soil eutrophication as well as elevated levels of temperature and CO2, that are associated with urban development (Thompson & McCarthy, 2008; Williams et al., 2015). Moreover, it has been observed that multiple species tend to exhibit an increased specific leaf area (SLA, the leaf area per unit mass) in urban areas in response to the enhanced soil nitrogen concentration (Knapp et al., 2009; Petersen et al., 2021), thus indicating a convergence in leaf properties in urban habitats (Lambrecht et al., 2016). It is evident that various plant species often display shared alterations in their phenotypic traits in reaction to ecological shifts linked with urbanisation. These findings indicate that urban surroundings bring about shared selective pressures on different species.

By contrast, species undergoing distribution reductions and expansions in urban environments may demonstrate different urban adaptations and responses. Generally, species that are restrictedly distributed in urban areas produce more seeds with low dispersal abilities as an adaptation to habitat fragmentation (Cody & Overton, 1996; Cheptou ey al., 2008). Previous research examining *Crepis* plants in the urban areas of Paris, France, has clearly demonstrated that urban populations contain a greater ratio of non-dispersive seeds without a pappus compared to suburban populations (Cheptou et al., 2008). A decline in seed dispersal ability in urban regions are proposed to be adaptation for surviving in restricted urban habitats (Cheptou et al., 2017; Dubois & Cheptou, 2017). Meanwhile, if there are species that have expanded their distribution in urban environments, they may have evolved more dispersive seeds compared to rural habitats. Thus, we can predict differing adaptative responses related to seed dispersal traits between species with restricted distribution and those that have expanded in urban habitat settings.

We aim to compare distribution and seed dispersal ability between two subspecies of *Youngia japonica*, both of which co-occur within the urban-rural gradient, to investigate the previously outlined predictions. *Youngia japonica* subsp. *Japonica* (hereafter, *Y.j. japonica*) and *Y*.*j*. subsp. *elstonii* (*Y.j*. *elstonii*) share most phenotypic traits and ecological characteristics (Shi et al., 2011; Fig. 1a,b). Meanwhile, a qualitative observation suggested that *Y.j. japonica* increases their dominance in urban areas whereas *Y.j*. *elstonii* shows decreased dominance in such environments (Shi et al., 2011). Nevertheless, the quantitative disparity in the distribution of both subspecies within urban-rural gradients has not yet been studied. At first, we qualitatively clarify whether *Y.j*. *japonica* and *Y.j*. *elstonii* undergoing distribution expansions and shrinkage in urban areas or not, respectively and environmental factors that cause the distribution difference between subspecies in a megacity area. Secondly, we examined changes in seed traits within the rural-urban gradient of this area in these two subspecies. Then, we analyse the interrelation between specific traits and the terminal velocity of each seed to understand whether each seed trait influences dispersal distance. Examining these contents, the following two questions are specifically addressed: (1) Are the distribution patterns of these two subspecies distinct in the urban-rural gradient, and can the difference be explained by environmental variations between urban and rural habitats? (2) Do urban populations of the subspecies with increased or decreased dominances in urban settings produce seeds with more or less dispersal ability compared to rural populations, respectively? Specifically, does *Y.j. japonica*, which is more dominant in urban areas, exhibits enhanced seed dispersal abilities whilst *Y.j. elstonii*, which is limited distribution in urban settings, showcase reduced seed dispersal abilities? Based on our results, we address these questions and discuss the relationship between distribution pattern and seed dispersal ability as urban adaptive responses in plants.

**Fig. 1:**
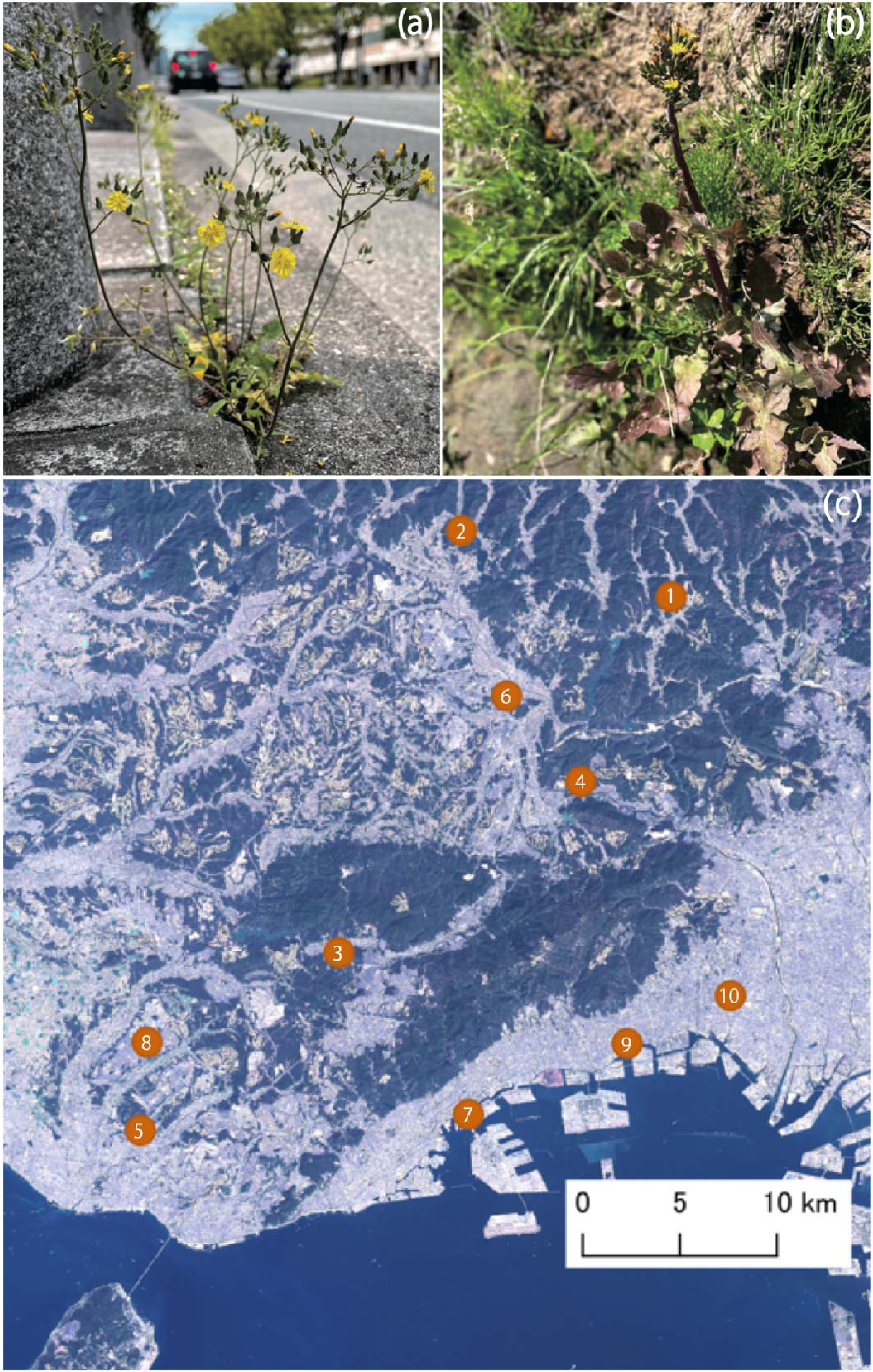
Photographs of main habitats of two subspecies of *Youngia japonica* and map: (a) *Yj. japonica* at the roadside; (b) *Yj*. *elstonii* at the paddy field; (c) Map of 10 study sites in the Osaka-Kobe metropolitan area. Site number shows order of urbanisation around study site (1000-m DLA) and lower number indicates lower levels of urbanisation. Sites 1∼6 are paddy field sites and sites 7∼10 are those in urban parks or roadsides (see Table S1)

## MATERIALS AND METHODS

### Study area and plots

The study was conducted 10 sites across the urban-rural landscape of the Osaka-Kobe metropolitan area (34°68′–96′ N, 134°01′–135°35′ E; Kobe, Sanda, Takarazuka and Nishinomiya cities, Hyogo prefecture). A survey plot (220 m square) was established at each of the 10 sites: six sites in rural and sub-urban paddy fields and four sites in urban parks or roadsides in the cities of Kobe, Takarazuka, Nishinomiya and Sanda (Fig. 1c, Table S1). In the study area, consolidated and unconsolidated traditional paddy fields, as well as neighboring secondary forests, are widely distributed in its rural areas (Uematsu et al. 2010), however, rapid urbanisation since the 1980s has led to the loss or fragmentation of plant habitats as rice fields and secondary forests are replaced by houses, buildings and paved roads (Tsuji et al. 2011; Uchida et al. 2018). All study paddy fields were found to have been maintained as paddy fields for at least 50 years and had not experienced any field consolidation, based on identification from historical aerial photographs (GSI maps). We selected paddy fields with semi-natural grasslands maintained by regular mowing on levees of paddies and irrigation ponds and forest edges (Uematsu & Ushimaru, 2013; Uchida & Ushimaru, 2014). The urban parks studied refer to parks surrounded by residential areas and green spaces within a campus of Kobe University, while roadside habitats were those on roadside green spaces. In Europe, such green spaces scattered throughout the urban areas have been reported to function as plant habitats (Jim & Liu, 2001; Stewart et al., 2009; Akinnifesi et al., 2010). The 10 sites were separated from one another by varying distances (Fig. 1c), with a minimum distance of 4.5 km.

### Environmental factors

For each study site, we calculated the total areas of developed lands (including residential/partly commercial, and industrial land uses, and areas for public facilities such as stations, city halls, and hospitals) within a 1 km radius from the site centre using QGIS software (version 3.16.1; QGIS Development Team 2021) and aerial images from Google Maps (2021). In this study, we calculated the total developed land area (DLA, ha) around each site at four different spatial scales (circular areas with radii of 250, 500, 750, and 1,000 m extending from each study site) as an index of degree of urbanisation for each site. In subsequent analyses, we presented the DLA at the radius that gave the lowest AIC value for the model.

### Study species

The two subspecies of *Y. japonica* (L.) DC. (*Y.j*. *japonica* and *Y.j*. *elstonii*) are annual to biennial herbs of the genus *Youngia* in the Asteraceae family and is found in the temperate regions of East Asia. The species is mainly found along the levees of rice fields and streams, forest edges, and roadsides (Fig. 1a,b). Both subspecies produce one to several stalk on a single plant, each containing about 1-40 flower heads with 5-25 ligulate florets (Shi et al., 2011; Ohashi et al., 2017). Each flower head buds are enclosed by involucres and open for a single day flower, typically from around 8am to midday. After flowering, the involucres close and the seeds with the pappus mature within them.

Two subspecies of *Y. japonica*, *Y.j. japonica* and *Y.j. elstonii* have been described based on differences in several vegetative and reproductive traits (Shi et al., 2011; Ohashi et al., 2017). The subspecies, *Y.j. japonica,* is characterised by green basal leaves still in the second year, a stem surface with few trichomes, multiple stalkless flowering stems and the production of flower heads of 8-13 mm in diameter (Shi et al., 2011; Ohashi et al., 2017). This subspecies tends to thrive in disturbed environments, such as roadside plantings, more often than *Y.j. elstonii* (Fig. 1a). In the study area, *Y.j. japonica* flowers from March to October. *Y.j. elstonii* is characterised by reddish rosset leaves in the second year, dense trichomes mainly on the lower part of the inflorescence stems, several relatively large stem leaves on each inflorescence stem and the production of flower heads with a diameter of 7-10 mm (Shi et al., 2011; Ohashi et al., 2017). Compared to *Y.j. japonica*, *Y.j. elstonii* is more commonly found in semi-natural ecosystems such as agricultural landscapes (Fig. 1b). In the study area, flowering of *Y.j. elstonii* occurs from April to August.

### Soil environmental factors

On the 5^th^ day of continuous rainfall absence, 26 July 2021 (with an average temperature of 28.7°C, a high of 33.8°C and a low of 25.5°C in Kobe), we investigated of soil environmental factors (soil moisture content and pH) in the habitats of the two subspecies of *Y. japonica*. We collected soil samples at a depth of 5 cm using a cylindrical soil corer with a diameter of 5-cm and a height of 5-cm (ca. 100 cm3) from four points, each located within a 30-m radius from the center of each survey plots. For each plot, the fresh soil samples weighted, dried at 40 L for 72 h, and weighted again. Soil water content (SWC in %) was calculated as follows (Nagata & Ushimaru 2016):

The average of the four samples at each site was used as the value representing that specific survey site.

In addition, soil samples passed through a sieve with a mesh size of approximately 2.4 mm were diluted with distilled water and the soil pH was measured using a pH meter (HI 99121; Hanna Instruments, Ltd., Woonsocket, RI, USA). Soil pH was determined by taking three measurements on each soil sample at each site and using the average value for analysis.

### Distribution of two subspecies within the urban-rural gradient

In April and May 2023, we investigated whether the distributions of *Y.j. japonica* and *Y.j. elstonii* differed according to the degree of urbanisation around their habitat. We counted the numbers of individual plants (abundance) of *Y.j. japonica* and *Y.j. elstonii* within each survey plot at each study site. We carefully surveyed accessible areas within each survey plot and identified subspecies of all *Y.j. japonica* individuals and recoded their locations. Wooded areas, water bodies and major roads in the study plots were excluded for survey areas due to their unsuitability for the growth of *Y. japonica* and safety concerns for the survey.

### Seed trait measurements

We collected seeds of both subspecies at each study site between April and June in 2021 and 2022 and measured seed traits including achene length, maximum achene diameter, and pappus length. Indicators such as plume loading (weight/projected area) and terminal velocity are commonly used to assess the dispersal ability of seeds with pappus. However, as *Y. japonica* seeds are too light to measure the mass of each achene, we calculated the seed ratio (pappus/achene length), which is considered to influence plume loading and terminal velocity and often used for an indicator of seed dispersal (Meyer & Carison, 2008). We collected seeds from 2-20 individuals of *Y.j. japonica* and 1-33 individuals of *Y.j. elstonii* at each study site. Owing to frequent mowing disturbances at the study sites, the sample sizes were notably restricted in some sites. We measured the seed traits for 10 seeds per individual (averaging 9.9 seeds) and used these measurements for our analysis. To measure the seed traits, we took images of the seeds using a scanner, calibrated the pixels using ImageJ software, and then recorded the measurements in mm. It should be noted, however, that at one site (paddy field, Shimosasori) in Takarazuka we could not collect *Y.j. japonica* seeds because the area had been mown before the complete seed sampling.

### Measurement of terminal velocities of individual seeds

To evaluate how seed trait values affect terminal velocity alterations, we gathered seeds from two *Y. japonica* subspecies across different study sites from October 2022 to April, May, and September 2023. We measured up to five seeds per plant (average of 4.9 seeds) to record their terminal velocity and seed traits which varied between urban and rural populations in either subspecies: achene length (mm), pappus length (mm), and seed ratio were measured. Although there were biases in the sampled seeds based on their subspecies and geography arising from the surveys’ timings, it is thought that the similarity of the seeds’ morphology from both subspecies negates any large effect of these biases on the terminal velocity calculation, from a physics-based point of view. Many samples were taken from urban *Y.j. japonica* individuals.

Terminal velocity is a vital factor in estimating dispersal ability of wind-dispersed seeds. Higher terminal velocity is considered to decrease seed dispersal distance (Andersen 1992; Zhu et al. 2016). An equipment to measure terminal velocity was developed based on Liu et al.’s (2021) research. After recording seed falling by a video camera (iPhone 12 mini, Apple, Cupertino, CA, USA), the UMATracker software (Yamanaka & Takeuchi, 2018) was used to track the seed coordinates. The camera’s perspective angle caused a discrepancy between the seed positions observed in three dimensions and those projected onto a two-dimensional scale. Therefore, in accordance with Liu et al.’s (2021) approach, we calibrated the position coordinates obtained from tracking to calculate the inter-frame movement speed. The apparatus configuration included a settling structure, standing at a height of 180 cm, and a camera with a black background. A scale with 1-mm precision was attached to the background, to facilitate calibration of the seed position. Acrylic shields and LED lights were placed outside the settling tower to enhance the clarity of images captured by the camera. An acrylic door was installed at the front of the settling tower to allow for recording seed movements. In addition, a seed release point was attached to the tower’s apex, which was positioned 180 cm above the ground, in order to minimize discrepancies in both horizontal and vertical seed falling positions. The iPhone was set at 30 cm from the backboard to recording the process of seed falling. The camera was set to record at a speed of 60 frames per second (fps). Considering the potential errors introduced during the tracking process, the average inter-frame movement speed for each seed was used as the terminal velocity in subsequent analyses.

Seed trait measurements were conducted after recording videos to mitigate the effects of pappus deformation during measurements.

## Statistical analyses

### Soil environmental gradients within the urban-rural gradient

To examine the effects of urbanisation in the surrounding areas on soil environmental factors, we constructed a general linear mixed model (LMM) in which soil pH or SWC was incorporated as the response variable, DLA of each spatial scale and a spatial autocovariate of the response variable were incorporated as the explanatory variables, the plot ID of each study site were incorporated as the random effect. The spatial autocovariate was calculated from latitude and longitude measurements of the centres of study sites to account for spatial autocorrelation effects on the analysis results (Dormann et al. 2007). We clarified the scale at which each soil factor was most affected by DLA using a model selection procedure based on Akaike’s Information Criterion (AIC).

### Distribution of two subspecies within the urban-rural gradient

To investigate the effects of urbanisation around the study sites or the soil environmental factors on the distribution (abundance or relative abundance) of the two subspecies, we constructed generalized linear mixed models (GLMMs) with Poisson distribution and log-link function for abundance data and with binomial distribution and logit-link function for relative frequency data, including the abundance or the relative abundance of each subspecies at different study sites as response variables, DLA of each spatial scale or the two soil environmental factors (SWC and pH) and the spatial autocovariate of response variable as explanatory variables. The scale at which each abundance metrics was most influenced by DLA was examined using an AIC-based model selection.

### Seed trait variations within the urban-rural gradient

To examine the effects of urbanisation around study site on seed traits, we constructed a LMM, where achene length, maximum achene diameter, pappus length, or seed ratio was incorporated as the response variable, DLA, subspecies identity (*Y.j. japonica*/*Y.j.elstonii*, 0/1 or 1/0) , the interaction between subspecies and DLA, and the spatial autocovariate were incorporated as the explanatory variables, the individual ID of seed mother and survey year were incorporated as random effect. In all analyses of seed traits, we analysed two models in which either of the two subspecies was used as a baseline in order to examine the significance of estimated coefficient of DLA for each subspecies. We examined the scale at which DLA most strongly influences each seed trait using an AIC-based model selection.

### Measurement of terminal velocities of individual seeds

To examine the effects of seed traits on the terminal velocity of *Y. japonica* seeds, we conducted a LMM. Each seed trait, namely achene length, pappus length, or seed ratio, in addition to subspecies identity (*Y.j. japonica*/*Y.j.elstonii*, 0/1) and the interaction between subspecies served as an explanatory variable, whereas terminal velocity was the response variable. The model included the individual ID of the seed parent as a random effect. We compared the models incorporating different seed traits as explanatory variables using the AIC, and the model that presented the best fit was chosen for further discussion.

All analyses in this study were performed using R software (version 4.1.2; R Core Team 2021). LMM and GLMM analyses were performed using the lme4, lmerTest and glmmTMB packages, and spatial autocovariates were calculated using spdep (Bivand et al., 2011; Bates et al., 2015; Brooks et al., 2017; Kuznetsova et al., 2017).

## Results

### Soil environmental gradients within the urban-rural gradient

The LMM analyses revealed that SWC and soil pH significantly decreased and increased with increasing the degree of urbanisation around the study site, respectively (Fig. 2, Table S3). Based on AIC value comparisons, models including DLA within a 750m and 250m radii from the study site centre were selected as the best models for the SWC and soil pH, respectively (Table S2).

**Fig. 2:**
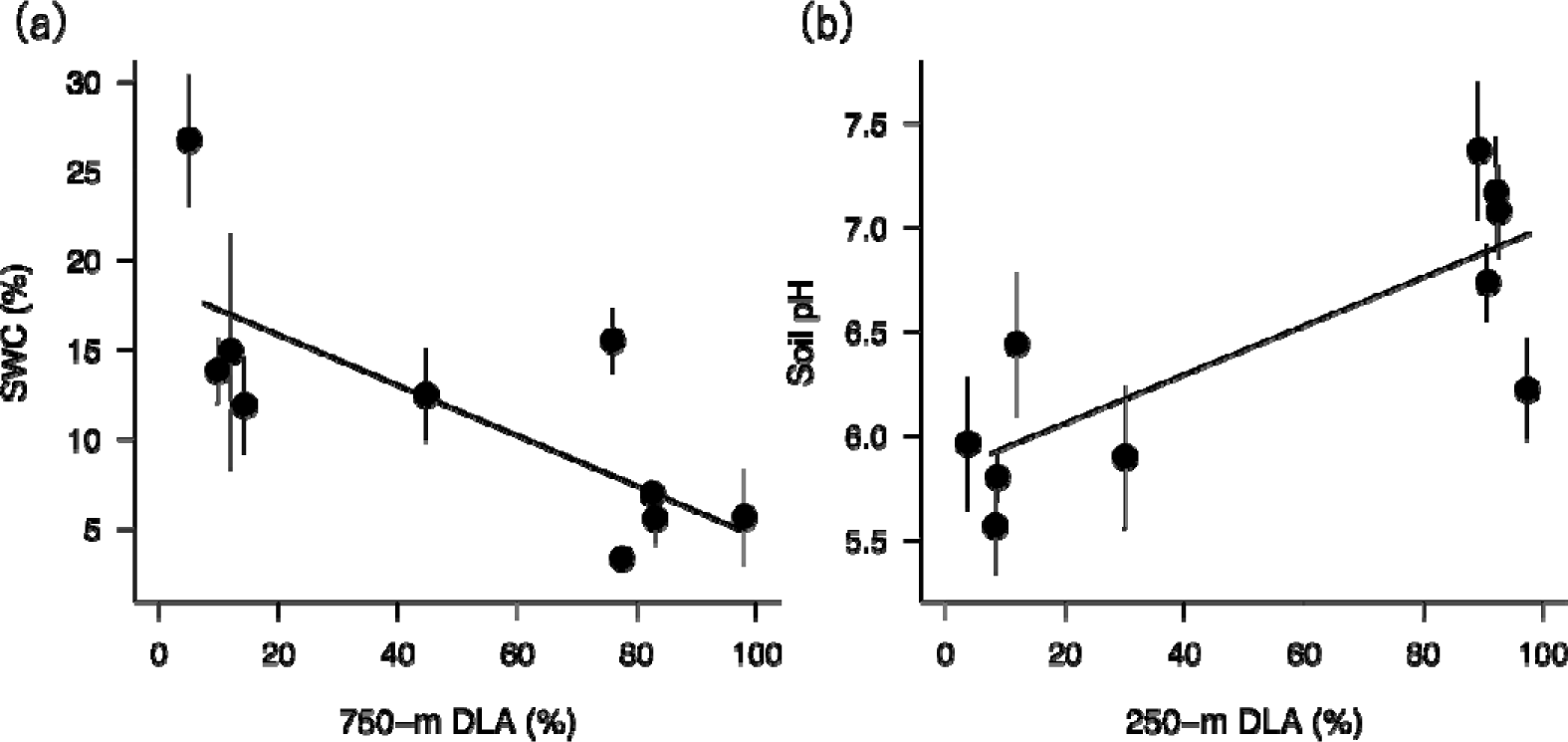
Relationships between the percentage of DLA of selected scale for each study site and soil environmental factors: (a) SWC on the 750-m DLA; (b) soil pH on the 250-m DLA. Each plot and error bar show mean and standard errors for each site, respectively. Regression lines were drawn using estimated coefficients from the general linear mixed models.

### Distribution of two subspecies along urban

In the distribution (abundance and relative abundance) analyses, models incorporating a 1000-m DLA were selected for both subspecies as the best models with the lowest AIC values (Table S4). The abundance of *Y.j. japonica* increased significantly with the degree of urbanisation (*p* < 0.001, Fig. 3a, Table S5), whereas the abundance of *Y.j. elstonii* decreased significantly (*p* < 0.001, Fig. 3a , Table S5). Significant relationships between soil pH and their abundance were observed for both subspecies. The number of *Y.j. japonica* and *Y.j. elstonii* individuals decreased and increased with increasing SWC, respectively (*p* < 0.001 for both subspecies, Fig. 3b, Table S5). Meanwhile, the number of *Y.j. japonica* and *Y.j. elstonii* individuals increased and decreased, respectively, with increasing soil pH (*p* < 0.001 and *p* < 0.001, respectively, Fig. 3c, Table S5). The pattern of the relative abundances of the two subspecies were similar to the absolute abundance patterns (Fig. 3d–f, Table S5).

**Fig. 3:**
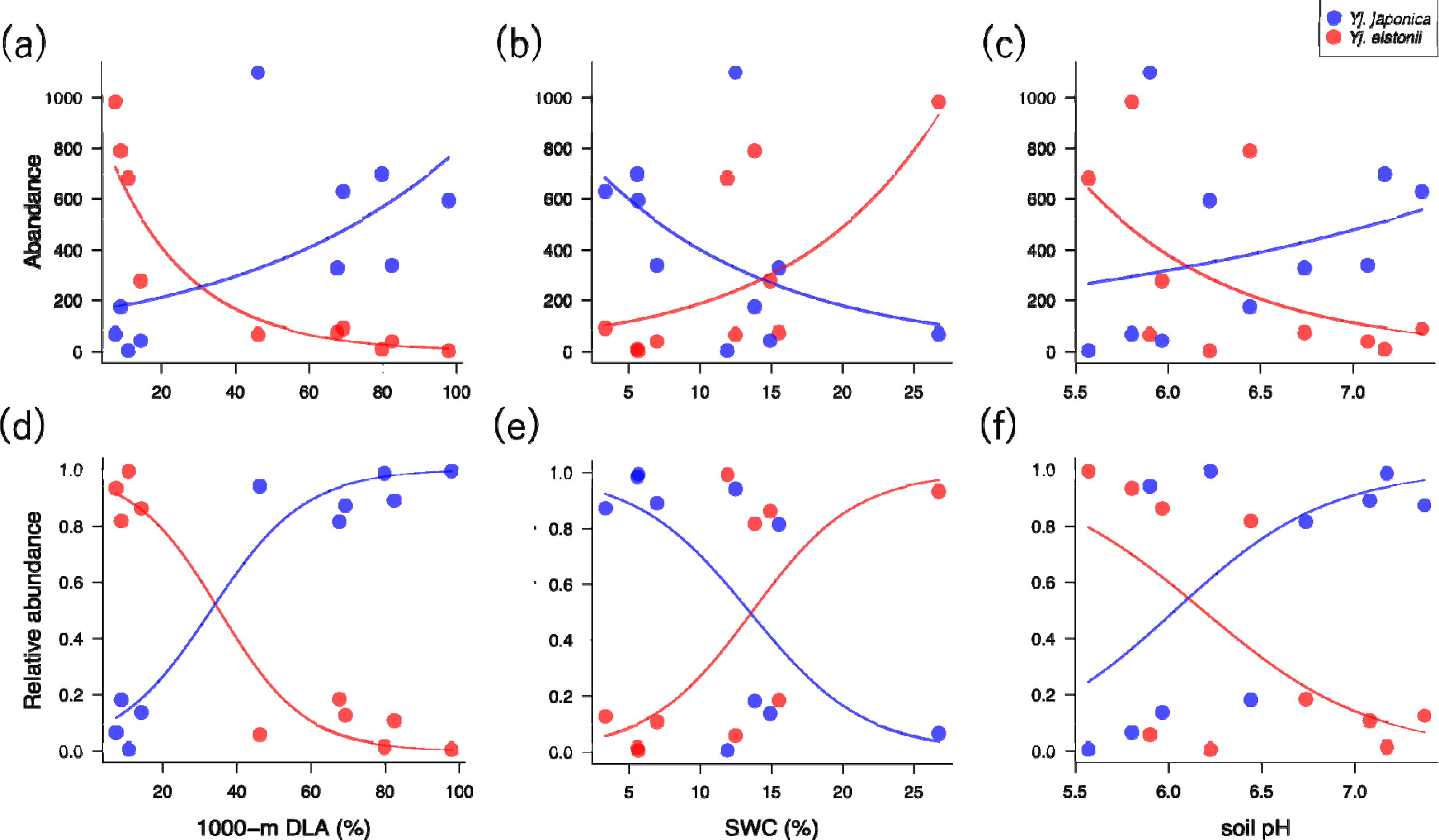
Relationships between the abundance (a–c) or relative abundance (e–f) of each subspecies of *Youngia japonica* and the degree of urbanisation and soil environmental facto d) 1000-m DLA (%); (b, e) SWC (%); (c, f) soil pH. Logistic regression lines were drawn using estimated coefficients from generalised linear models with Poisson distribution and those with binomial distribution (d–f). Colours indicate subspecies; blue, *Yj. japonica*; red, *Yj. elstonii*.

### Seed trait variations within the urban-rural gradient

In all the analyses of seed traits, models incorporating a 250 m DLA was selected as the best models with the lowest AIC values (Table S6). Both subspecies exhibited a decrease in the seed ratio with increasing urbanisation (*Y.j. japonica*, *p* < 0.05; *Y.j. elstonii*, *p* < 0.001, Fig. 4a, Table S7), although the factors causing urban-rural variations in the seed ratio differed between the subspecies. Achene length significantly increased with increasing the degree of urbanisation in *Y.j. japonica* (*p* < 0.01, Fig. 4b, Table S7), whereas no significant relationship between them was observed in *Y.j. elstonii* (*p* = 0.105, Fig. 4b, Table S7). On the other hand, there was no significant relationship between pappus length and the degree of urbanisation in *Y.j. japonica* (*p* = 0.445, Fig. 4c, Table S7), while in *Y.j. elstonii* pappus length significantly decreased with increasing the degree of urbanisation (*p* < 0.001, Fig. 4c, Table S7). Regarding maximum achene diameter, neither subspecies showed significant variation in response to urbanisation around the study site (*Y.j. japonica*, *p* = 0.578; *Y.j. elstonii*, *p* = 0.215, Fig. 4d, Table S7).

**Fig. 4:**
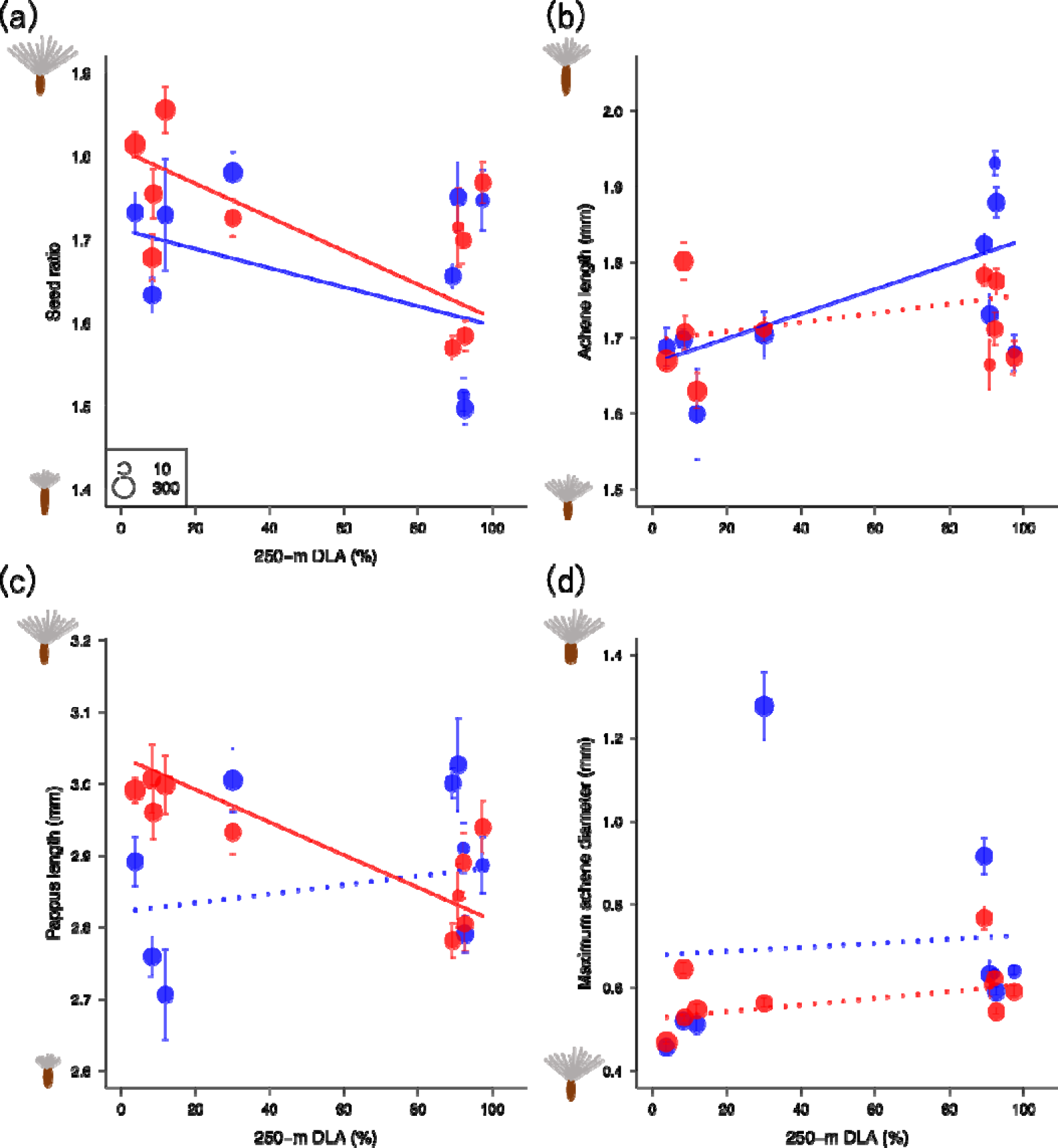
Relationships between the percentage of 250-m DLA of each study site and seed traits: (a) seed ratio ; (b) achene length (mm); (c) pappus length (mm); (d) maximum achene length (mm). Each plot and error bar show mean and standard errors for each site, respectively. Plot size expresses sample size for each site, respectively. Regression lines were drawn using significant estimated coefficients from the general linear mixed models. Dotted lines indicate non-significant relationships. The colors indicate subspecies; blue, *Yj. japonica*; red, *Yj. elstonii*.

### Measurement of terminal velocities of individual seeds

Based on the AIC value comparison, the model including pappus length, species identity and their interaction as the explanatory variables was selected as the best fit model (Table S8). The terminal velocities of individual seeds significantly decreased with increasing their pappus length (Fig. 5, Table S9), Meanwhile, species identity and the interaction with pappus length had no significant effects on terminal velocity in the model (Table S9). It should be noted that seed ratio and achene length had no effect on it in the LMMs (Fig. S1, Table S9).

**Fig. 5:**
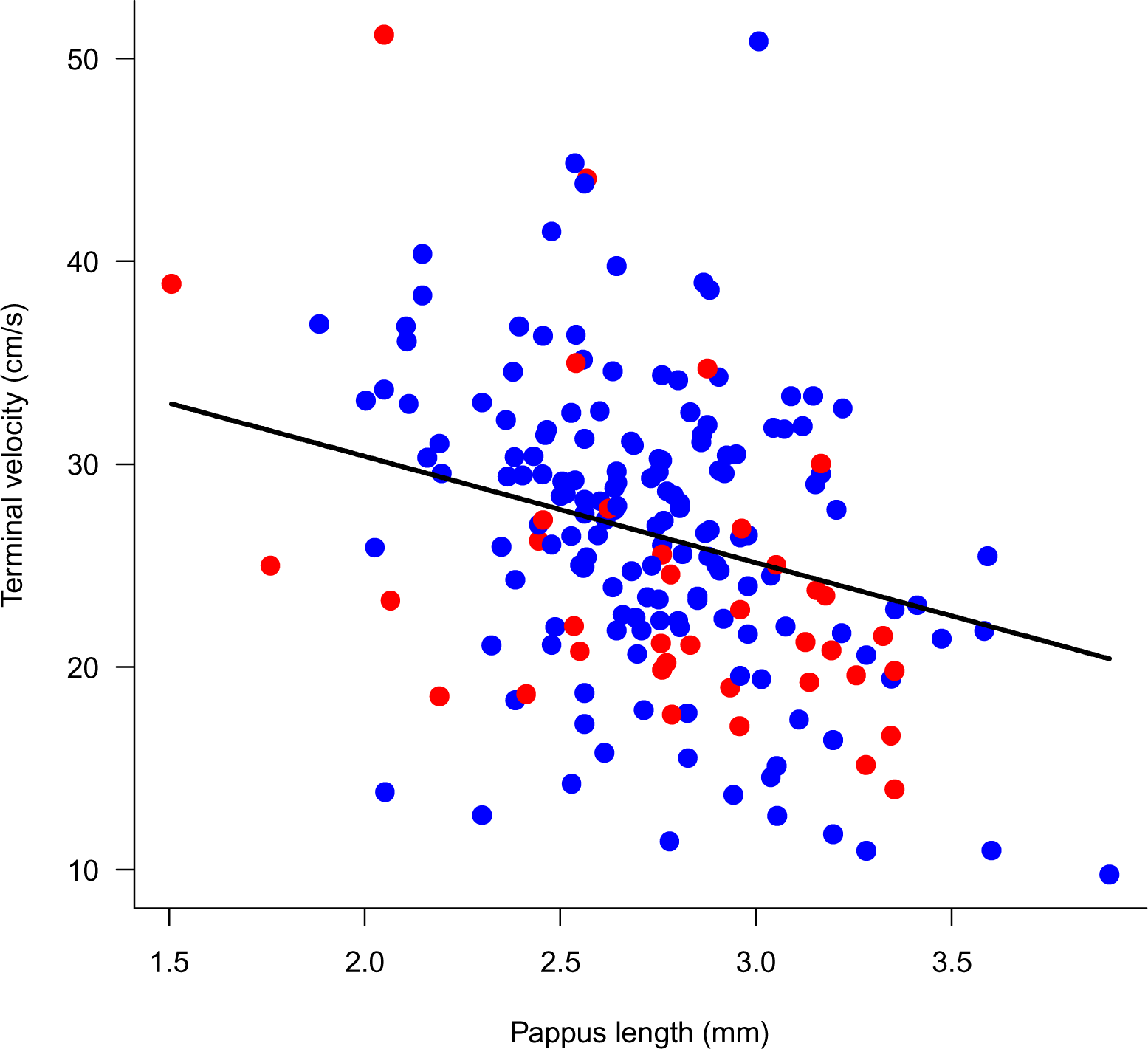
Relationships between the pappus length and terminal velocity. Each plot shows terminal velocity and pappus length for each seed. A regression line was drawn using significant estimated coefficients from the general linear mixed models. The colors indicate subspecies; blue, *Yj. japonica*; red, *Yj. elstonii*.

## Discussion

The two subspecies of *Y. japonica* were distributed differently within the urban–rural gradient of the Osaka-Kobe megacity. Our finding suggests that *Y.j. japonica* thrived more in drier and neutral soil conditions in urban areas, while *Y.j. elstonii* dominated more in moister and more acidic soil conditions of rural areas. In contrast to our expectations, urban populations of both subspecies had significant lower seed ratio than their rural counterparts. Lower pappus length caused higher terminal velocity, indicting lower dispersal abilities in urban *Y.j. japonica* individuals compared to their rural counterparts. More detailed discussions on our findings are made below.

### Differences in distribution between two subspecies within urban-rural gradient

This study newly provides quantitative evidence for changes in the absolute and relative abundance of the two subspecies of *Y. japonica* along the urban-rural gradient, supporting previous qualitative observations (Shi et al., 2011). Furthermore, we revealed that changes in SWC and soil pH along the urban–rural gradient might be key factors explaining the abundance change of each subspecies. The findings support previous European studies showing that urban environments are experiencing soil drying and increased soil pH (Pickett et al., 2011; Asabere et al., 2018), suggesting that habitat environmental changes due to increased artificial land cover are occurring in Asian megacities in a similar way to European cities and have significant impacts on plant distributions. In urban habitats with drier soil conditions with near-neutral pH, *Y.j. elstonii* shrank the population sizes whereas *Y.j. japonica* expanded the distribution. Niche partitioning between closely related herbaceous plant species due to differences in soil drought stress tolerance is reported in *Mimulus guttatus* and *M. laciniatus*. (DeMarche et al., 2013). The two subspecies of *Y. japonica* are usually sympatrically distributed in rural areas as our study demonstrated (Fig. S2). To reduce competition between the subspecies, niche partitioning along micro soil environmental conditions might occur, likely causing the difference in response to newly created urban soil conditions.

### Urban–rural variations in seed traits

Urban individuals of both subspecies commonly exhibited decreased seed ratios independent of their dominances in urban areas. Although the reduction in seed ratio is predicted to decrease seed dispersal capability (Meyer & Carison, 2008), no significant relationship was found between terminal velocity and seed ratio.

By examining the individual contributions of achene length and pappus length to the seed ratio, different patterns of response to increasing urbanisation are observed in both subspecies. In *Y.j. elstonii*, reduced pappus length in urban populations contributes to the reduction in seed ratio. As we found the significant negative relationship between the length of the pappus and the terminal velocity, a reduction in the diameter of the parachute portion owing to the shorter pappus may directly reduce seed dispersal distance in this subspecies (Fig. 5; Meyer & Carison, 2008). Thus, the decrease in pappus length likely reduces seed dispersal abilities in urban *Y.j. elstonii* populations compared to their rural counterparts.

On the other hand, a decrease in pappus length was not found in urban *Y.j. japonica* populations, instead an increase in achene length contributes to the decrease in their seed ratio. Increased achene length suggests an increase in seed size. In plants, increased seed size often increases germination rate (van Mölken et al., 2005; Cappuccino et al., 2002) and allows for earlier seedling growth (Meyer & Carison, 2008). These knowledges lead us to the hypothesis that *Y.j. japonica* may be more competitive in the early stages of life cycle in urban habitats than in rural habitats. Previous studies proposed the idea that urban plants would evolve earlier phenology in response to irregular harsh environments such as soil temperature raise and drying associated with urbanisation (Fitter & Fitter, 2002). In urban *Y.j. japonica* populations, increased seed size may allow early growth, which is adaptative in the urban dry habitat conditions. Increasing achene volume is costly in terms of dispersal capacity because increased achene volume rapidly increases seed fall rate due to increased pappus load (Greene & Johnson, 1993; Meyer & Carison, 2008). However, the increased achene length of urban *Y.j. japonica* may not be a major cost in dispersal ability, which was indicated no significant relationship between achene length and terminal velocity in the measured seeds (Fig. S1).

As an alternative explanation for the increase in seed size in urban *Y.j. japonica* populations, it could be the results of enhanced outcrossing. Increased outcrossing in angiosperms can lead to increased male-male competition and potentially increase seed size (Raunsgard et al., 2018; Larios & Mazer, 2022). Increased abundance (population size) of urban *Y.j. japonica* indicates an increase in potential mating partners. In addition, we observed frequent pollinator visits in urban populations of *Y.j. japonica* (Nakano & Ushimaru, unpublished data). Abundant mating partners and pollinators can maintain higher outcrossing rates, which may lead to larger seed size in urban *Y.j. japonica* individuals. In addition, seed size can also be influenced by the nutrient conditions of the parent plant (Larios & Mazer, 2022). It would therefore be important for future research to assess the validity of these ideas by investigating whether the outcrossing rate and soil nutrient conditions on seed size in *Y.j. japonica*. In addition, for *Y.j. japonica*, it would be necessary to determine the extent to which increased seed size contributes to seed survival and early growth in urban environments.

In this study, our finding that pappus length can be used as an indicator of terminal velocity offers valuable insights, especially for estimating the dispersal ability of small-seeded wind-dispersed plant species whose plume loadings are impossible to calculate due to the difficulty in measuring their seed weight. Because wind-dispersed herbs dominated in urban areas (Hayasaka et al., 2011), the seed ratio would be effective metrics to estimated their dispersal ability.

### Different responses to urban environments between the two subspecies

This study firstly compare the seed characteristics for the urban adaptation between closely related plant species that are expanding and shrinking their distributions in urban areas. Our findings highlight the potential differences in seed adaptation to urban environments. The results of this study suggest that in urban environments, where many plant species are experiencing reduced population sizes owing to habitat fragmentation, there is potential for some species to expand their distribution through trait adaptation. Additionally, in wind-dispersed plants whose seeds are too small to measure their weights, the present study suggests that pappus length could be a usefull indicator of seed dispersal ability. It is important to note that we investigated only a single pair of closely-related species in a single megacity area. Therefore, future research should investigate the differences in urban adaptation between other closely-related species pairs in various urban areas of different geographical regions to test our hypothesis and findings.

## Acknowledgements

We thank T. Saga for teaching us UMATracker and Y. Uchiyama for allow us using Arc GIS pro. We are grateful to paddy landowners who allowed our fieldwork on their lands. We also thank members of Biodiversity Laboratory, Minamoto Laboratory and Evolutionary Ecology Laboratory at Kobe University for giving us valuable comments on our study.

## Author contributions

SHN and AU conceived the study, designed the methodology; SHN collected all the data; SHN and AU analysed the data and wrote the manuscript.

