## Supplementary material for "Contrasting urban adaptations of seed dispersal traits in two closely-related plant species": Fig. S1

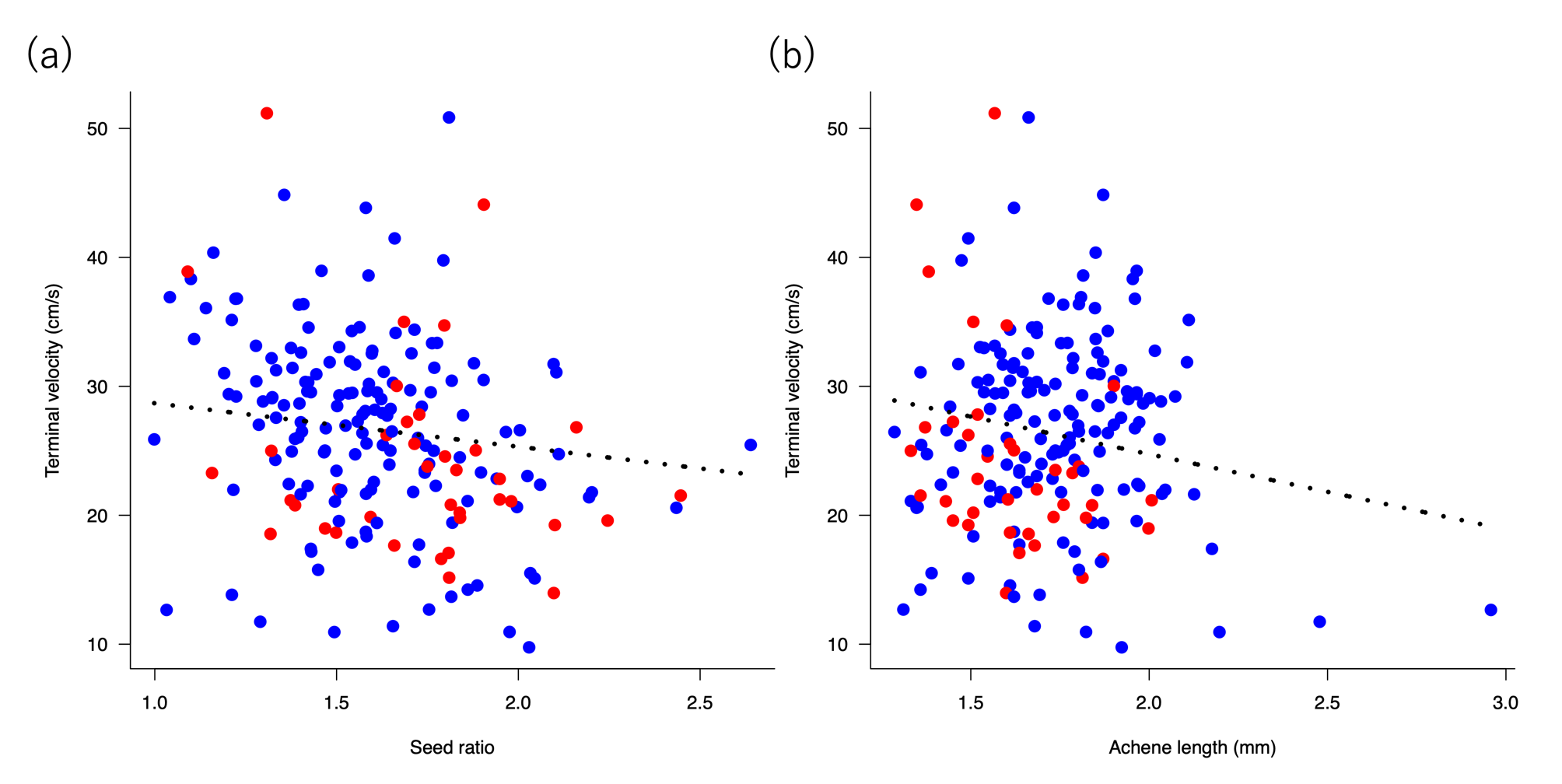

Fig. S1: Relationships between the seed ratio and achene length and terminal velocity. Each plot shows terminal velocity and seed traits for each seed, respectively. Dotted lines indicate non-significant relationships from the general linear mixed models. The colors indicate subspecies; blue, *Yj. japonica*; red, *Yj.elstonii*.
