## Supplementary material for "Contrasting urban adaptations of seed dispersal traits in two closely-related plant species": Fig. S2

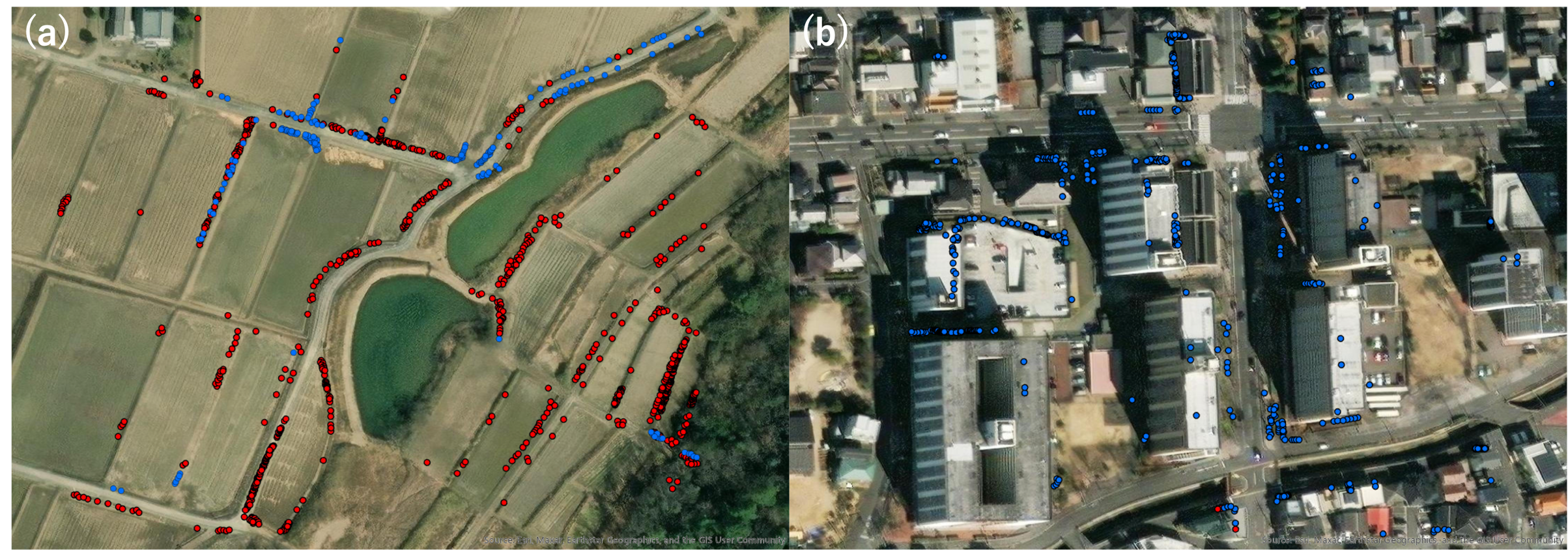

Fig. S2: The figures show the sympatric distribution of two subspecies in a rural study site (a, paddy fields, Sumada) and an urban study site (b, roadside, Nishinomiya). Blue and red dots indicate the distribution of *Y.j. japonica* and *Y.j. elstonii*, individuals, respectively. These photos create from using the Arc GIS.
